# Phloem conductance from leaf to fruits in avocado during development

**DOI:** 10.64898/2026.10.31.762750

**Authors:** María Agulló-Samper, Cristina Ferrer-Blanco, Carlos E. González, Sabrina Gennis, Kaare H. Jensen, Juan M. Losada

**Affiliations:** Institute for Mediterranean and Subtropical Horticulture ‘La Mayora’ (IHSM LaMayora-CSIC-UMA). Avda. Dr. Wienberg s/n. Algarrobo-Costa, 29750, Málaga, Spain; Department of Physics, Technical University of Denmark, DK-2800 Kgs. Lyngby, Denmark

**Keywords:** Phloem, sinks, sources, modelling, sieve tube, avocado, fruits

## Abstract

Reproductive function in seed plants depends on efficiently allocating nutrients from sources to developing fruits. Hydraulic transport of carbohydrates largely depends on phloem architecture, but how phloem structure in fruit pedicels changes with development remains unexplored for most seed plants. Using the one-year phenology of avocado trees, we quantified water content, dry matter, and non-structural carbohydrates (NSCs) in mature leaves, flowers, young fruits, and fully developed fruits. Concomitantly, we measured phloem conduit geometry in different vein orders of leaves and in flower-fruit pedicels over time, including sieve tube numbers in cross sections. NSCs were below 5% in leaves and fruits but reached the highest concentrations in flowers, especially fructose (20%) and glucose (35%), and the pedicels of flowers showed the smallest conduits. While sieve tube elements in leaves scaled their geometry from the minor veins to the petioles, in fruit pedicels, their geometry changed only from flower to initial fruit set, increasing their number linearly up to fruit maturation. Our models show that phloem transport toward the flowers is impaired, but total conductance increases gradually due to phloemogenesis.

## INTRODUCTION

The structure of the phloem in source and sink organs plays a fundamental role in resource allocation in plants during development, influencing key processes such as reproductive function. However, research on comparing the structure of the phloem in sources and sinks remains unstudied in most vascular plants. The mass flux of photo-assimilate transport and delivery are critically influenced by the geometry of the conduits transporting them, which includes length, radius, sieve plate pore size and number, and sieve tube number in cross-section (members of our team; Horwitz, 1954; Thompson and Holbrook, 2003; Jensen *et al*., 2013). The scaling geometry of the individual conduits that form a continuum in the phloem, or sieve tube elements (STEs), determines their hydraulic efficiency (Jensen *et al*., 2012ab; Savage *et al*., 2017; Barceló-Anguiano *et al*., 2021b), but studies focused on the geometry of the phloem conduits in the different vein orders have only recently emerged. So far, phloem conduit scaling in leaves has been studied in a limited number of species that include different growth forms, such as lianas (Losada *et al*., 2019), shrubs (Losada & Holbrook, 2019), and trees (Carvalho *et al*., 2018; Barceló-Anguiano *et al*., 2021a,b; He *et al*, 2024). All these works converge on the scaling relationship between vein order and sieve tube element size. Yet, the number of conduits, which plays a critical role in the bulk mass flow at the organ level, remains unexplored in most species with reticulate venation, except for leaves with a single vein, such as the needles of pines (Rosenberg *et al*., 2022; see also Vincent *et al*., 2025). The lack of information regarding the number of conduits in other species limits our understanding of net photo-assimilate transport from source to sink organs.

Developmentally, the mature leaf reached the maximum differentiation of vascular bundles; thus, it can be considered static (Esau, 1969). In contrast, the continuous gain of matter during fruit development putatively requires differentiation of the vascular cambium into more conductive phloem, but whether the number of conduits changes in fruit pedicels over time remains unexplored. At the same time, vascular flow rates have been measured mainly in the xylem of fruit pedicels, revealing a direct relationship with dry matter accumulation in several fruit trees, like kiwifruit (Clearwater *et al*., 2012; Morandi *et al*., 2010), grape (Greenspan *et al*., 1994), apple (Lang, 1980), peach (Morandi *et al*., 2007), pear (Morandi *et al*., 2014), sweet cherry (Brüggenwirth *et al*., 2016; Winkler *et al*., 2016), and avocado (Kaneko *et al*.,; 2024). Yet, these studies focus on functional outcomes (water/flow rates) rather than the underlying structural constraints of the phloem conduits (size and number) that ultimately determine the long-term hydraulic capacity for resource allocation.

While water drives carbohydrates towards fruits, they can either be stored, as in grapes, or transformed into lipids, as in avocados. Carbohydrates, synthesized in the foliar cells of the mesophyll, are loaded into the phloem conduits of the minor veins, increasing their osmotic pressure, which allows the passive entrance of water from the xylem, and that initiates a flow through the central veins, the petiole, the branches, eventually unloading in the growing organs, where sugar unloading occurs, dropping the osmotic pressure and thus allowing for directional transport (Münch, 1930). While daily changes in carbohydrate transport are well described (Brauner *et al*., 2018), little is known about seasonal changes that govern fructification. The rate of vascular flow changes seasonally and is constrained by the architectural construction of the phloem conduits (size and number) over time (Vincent *et al*., 2025). However, only a few studies have examined the structure and function of the phloem in developing fruits (Savage *et al*., 2015), and no quantification of the individual phloem elements in developing fruits has been reported to date. This is because distinguishing the sieve tube elements of the phloem requires a combination of microscopy and immunocytochemistry (Torode *et al*., 2018), which are typically costly and time-consuming. Comparing the morphology and number of sieve tube elements between petioles (sources) and pedicels (sinks) is essential for understanding the critical developmental stages of plant reproduction, with implications for ecology and agriculture. This research aims to shed light on how the vascular system of avocado pedicels changes over time during fruit growth, compared with that of petioles.

Avocado is increasingly becoming a staple food worldwide, and its cultivation has been rising by more than 100% in the last decade, with Mexico as the top producer. In Continental Europe, where the market of avocado grew 16% in 2025, Spain is the major producer, centered in the Mediterranean coast of the south, but expanding to other coastal areas (Rodríguez Pleguezuelo *et al*., 2018). It is also one of the highest latitudes for avocado cultivation, which influences their productivity rates, with one fruit per year compared with the two- or three-sometimes fruit production in tropical areas of the world (López-Hoyos *et al*., 2024). Due to its growing economic and social impact, numerous studies have focused on breeding and agronomic improvement of avocado fruits, but challenges of climate change require a physiological approach. Recent physiological studies on avocado focused on the hydraulics of the xylem under drought (Cardoso *et al*., 2020) or on the evaluation of vascular flows toward the fruits (Kaneko *et al*., 2022). Yet, there are no reports on the anatomical structure of avocado pedicels, even though it is crucial to understand the hydraulics of sugar transport to growing fruits. We used the most widely consumed variety worldwide, ‘Has’, to evaluate non-structural carbohydrates in leaves and fruits over time, and to compare phloem structure in leaf petioles and fruit pedicels. Our results reveal that phloem differentiation in fruits plays a critical part in the hydraulic transport of nutrients toward developing fruits.

## RESULTS

### Phenology of avocado cv. ‘Has’ vegetative and reproductive organs

The developmental cycle of avocado trees in the subtropical south of Spain takes about 12 months (**Figure 1**; also described in Alcaraz *et al*., 2013). Following the mild winter, almost all leaves drop off, concomitant with the emergence of inflorescences around March-April (**Figure 1A**). During the pollination period, inflorescences containing tens of tiny yellow flowers at the tips of the branches (hundreds per tree) coexist with a few developing leaves of the current year (**Figures 1B-C**). Following pollination, a small proportion of the flowers set fruit (Cameron *et al*. 1952), and reached the maximum percentage of water (80%) and around 18% of dry matter (Fig S1). Concomitantly, leaves of the first flush extend over a period of approximately 30 days (until the end of May), when they reach a complete expansion (**Figures 1D-F**). Fruits took four months (October, month 10) to reach complete expansion, and two months later (month 12), they reached full maturity and displayed a reduction in water content to 70% and an increase in dry matter to 28% (**Supplemental Figure S1**). Water percentage was minimal in flowers (10%), contrasting with the 70% in leaves and developing fruits, which reached a maximum of water during the initial stages of fruit set (**Figure1G**).

**Figure 1.**
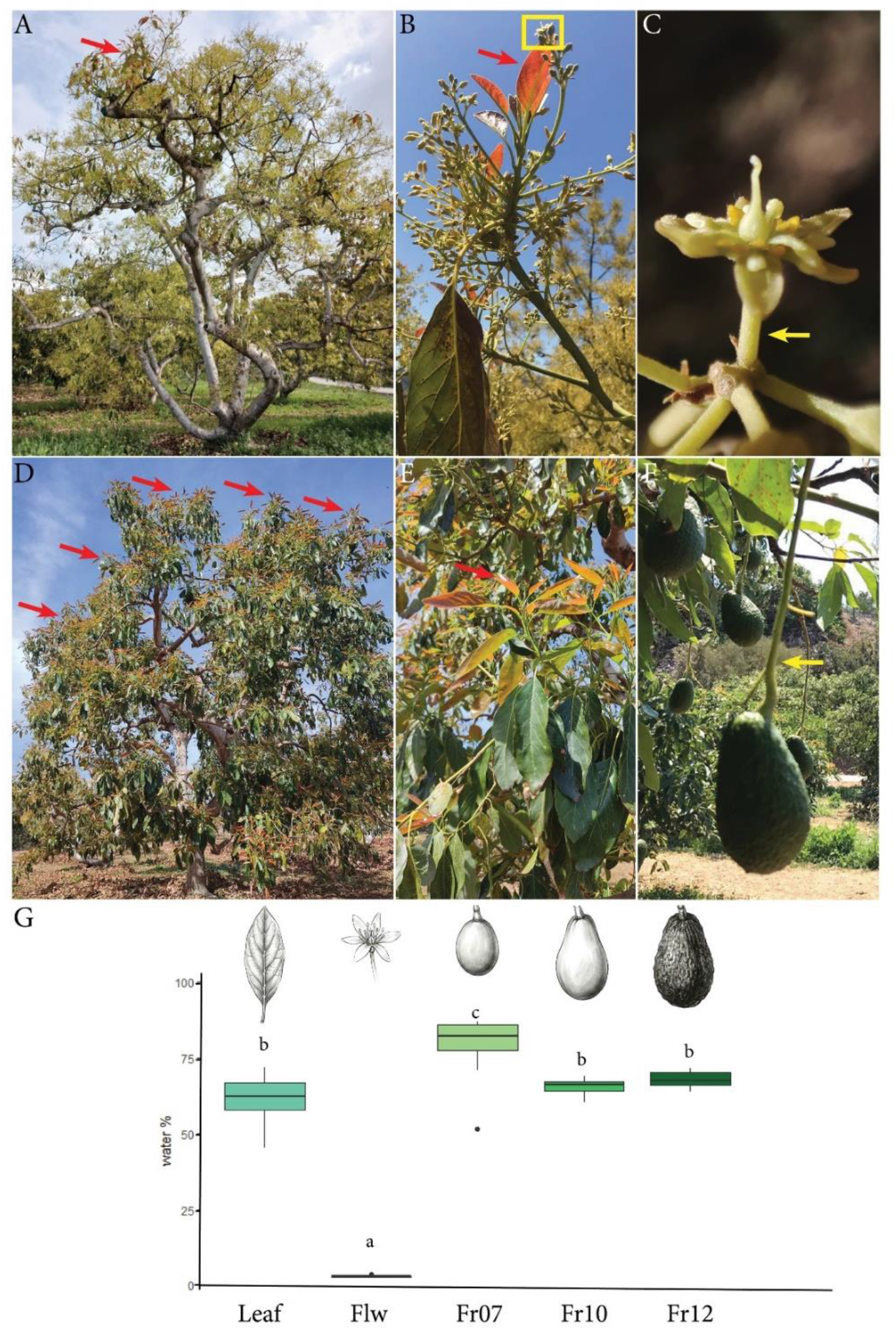
Morphology of mature avocado trees during flowering and fruit development. (A) Avocado tree at the stage of full bloom showing mostly inflorescences. (B) Detail of an inflorescence with terminal new vegetative growth (red arrow). (C) Individual flower at anthesis subtended by the pedicel (yellow arrow). (D) Mature tree during fruit growth covered with leaves and showing a second flush of leaves (red arrows). (E) Detailed view of the second flush of leaves (red arrows). (F) Developing avocado pending on the pedicel (yellow arrow). (G) Water content (%) of leaves, flowers, and fruits at different developmental stages, represented by the month of growth (7: July; 10: October; 12: December). Letters over bars indicate significant differences (ANOVA and Tukey’s post hoc test, *P* < 0.05).

The length-to-width ratio of reproductive organs increased from below 1.5 in flowers to above this value during the initial fruit set (**Figure 2A**). Still, it remained between 1.5 and 2.0 from the fruitlets to the mature fruits, without significant differences. Similarly, the diameter of the flower pedicels was 2mm (**Figure 2B**), and increased significantly at the initial fruit set in July, reaching a maximum of 5mm. Still, this value remained unchanged for months, until the fruits reached maximum expansion in October and subsequently matured in December.

**Figure 2.**
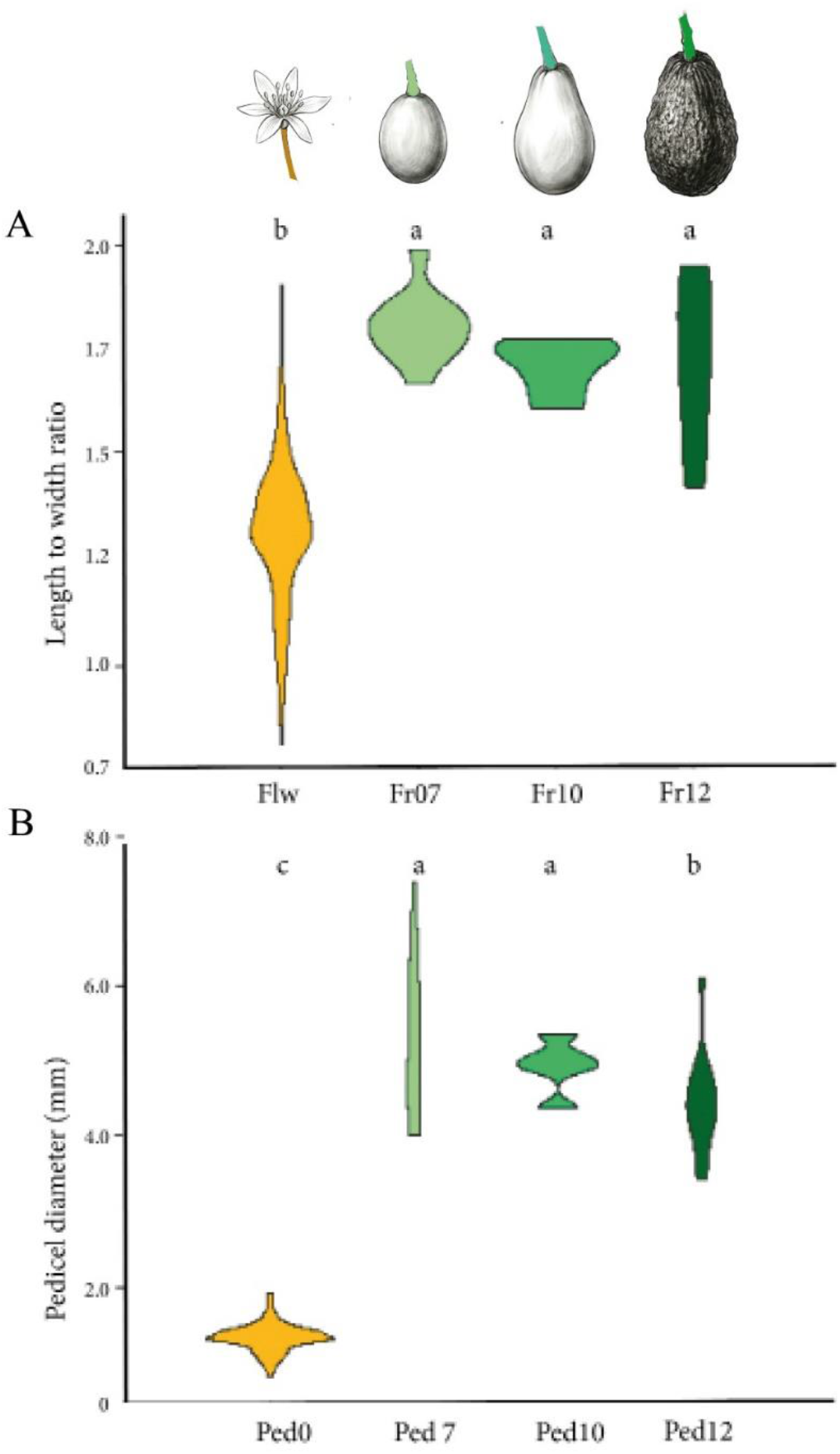
Fruit and pedicel grow during development. (A) Lenth to width ratio of the flower ovary (orange, Flw) and the developing fruits at months 7 (initial fruit set, pale green), 10 (maximum size attained, light green), and 12 (maturity, dark green). (B) Pedicel diameter at the different developmental stages. Letters over bars indicate significant differences (ANOVA and Tukey’s post hoc test, *P* < 0.05).

### Non-structural carbohydrates in avocado leaves and fruits during development

Comparison of starch in fully developed leaves from the previous and the current year revealed a similar amount of starch, around 5% (**Figure 3A**). Flowers reached a peak of ∼9% of starch, which was later reduced significantly to 2% in developing fruits, reaching a minimum of 0.5% in the mature fruits. Sucrose content was below 5% in all organs and fruit developmental stages, with little differences among them (**Figure 3B**). The content of fructose and glucose followed similar patterns, with 5% or lower in the leaves, a maximum amount in the flowers (20% and 35% w/w respectively), which decreased to the developing fruits, reaching a minimum of ∼1% in the mature fruits (**Figures 3C, D**).

**Figure 3.**
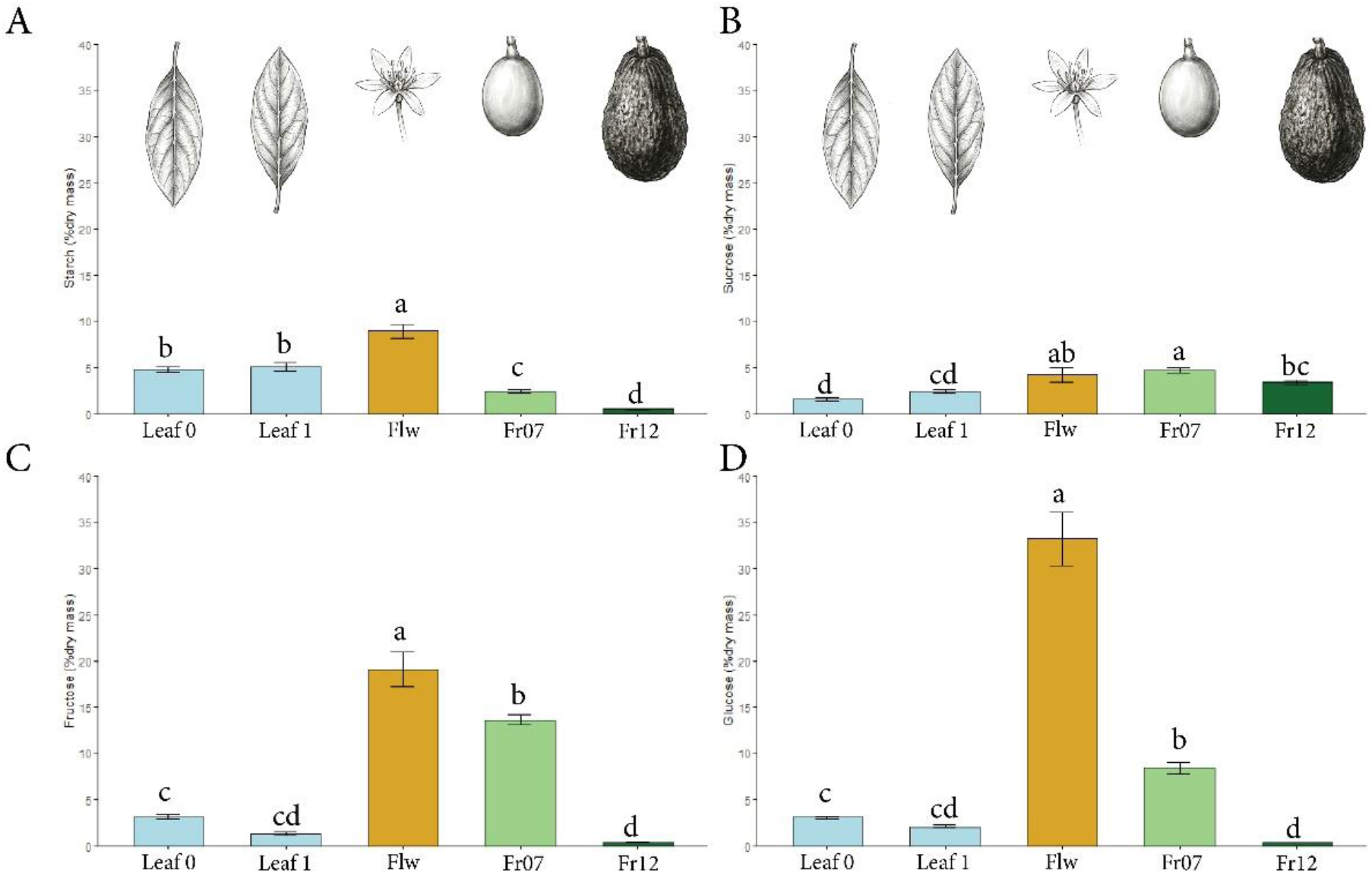
Non-structural carbohydrate (NSC) content in avocado leaves, flowers, and developing fruits. Concentrations are shown for leaves of the previous year (Leaf 0, outline pointing down), the current year (Leaf 1, outline pointing upright), flowers (Flw), and developing fruits in July (Fr07) and December (Fr12). (A) Starch. (B) Sucrose. (C) Fructose. (D) Glucose. Letters over bars indicate significant differences (ANOVA and Tukey’s post hoc test, *P* < 0.05).

### Phloem anatomy in the petioles and pedicels of developing avocado fruits

#### Length and width

Evaluations of the sieve tube elements in the different vein orders of the avocado leaves (**Figure 4A**) revealed a scaling relationship that was in line with the vein size (i.e. vein order). Thus, the minor veins showed the shortest and thinnest sieve tube elements, which increased in length to the veins or higher orders (**Figures 4B-C**). The STE length in the petiole was smaller than that in the primary vein, and similar to the secondary order veins. Similarly, the width of the sieve tube elements increased gradually from minor to major veins, and the sieve tubes of the petiole and the major veins were similar in diameter.

**Figure 4.**
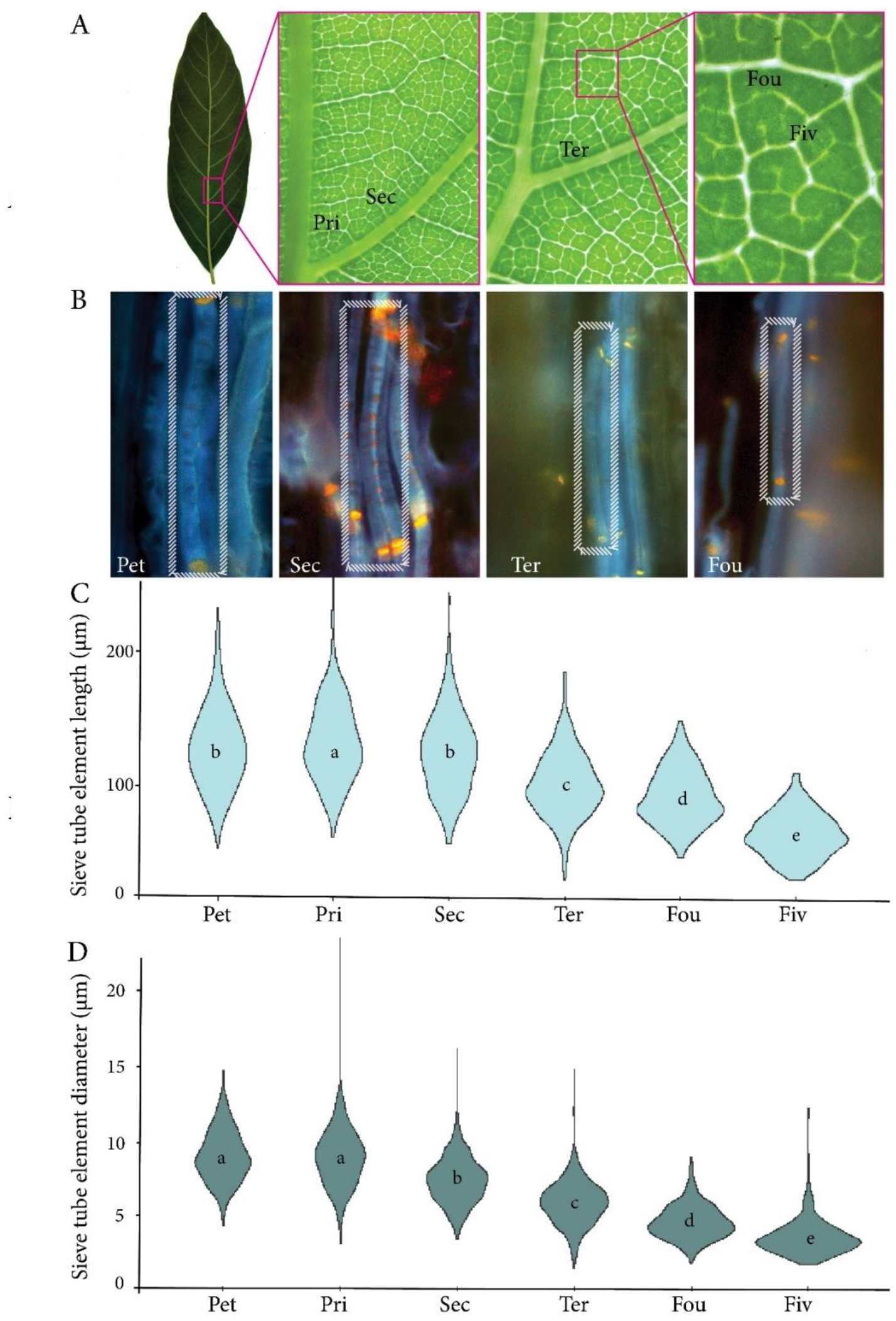
Phloem geometry across leaf vein orders in avocado. (A) Avocado leaf and close-ups of the different vein orders starting in the petiole (Pet), primary vein (Pri), secondary vein (Sec), tertiary vein (Ter), fourth-order veins (Fou), and fifth-order veins (Fiv). (B) Fluorescence images of individual sieve tube elements (dashed rectangle) stained with aniline blue for callose (yellow) and counterstained with Calcofluor White for cellulose (blue) at the indicated vein orders. (C) Sieve tube element length per vein order. (D) Sieve tube element diameter per vein order. Letters within violin plots indicate significant differences (ANOVA and Tukey’s post hoc test, *P* < 0.05).

Measurements of the phloem in the pedicels of reproductive organs revealed that the smallest sieve tube element width corresponded with the pedicels of flowers, increasing slightly to developing fruits (July or month 7 of development), maintaining the same values until maturity in December (**Figure 5A**). The length of the sieve tube elements was also minimum at the flower stage, and increased some microns significantly from flower to fruit development and maturity (**Figure 5B**).

**Figure 5.**
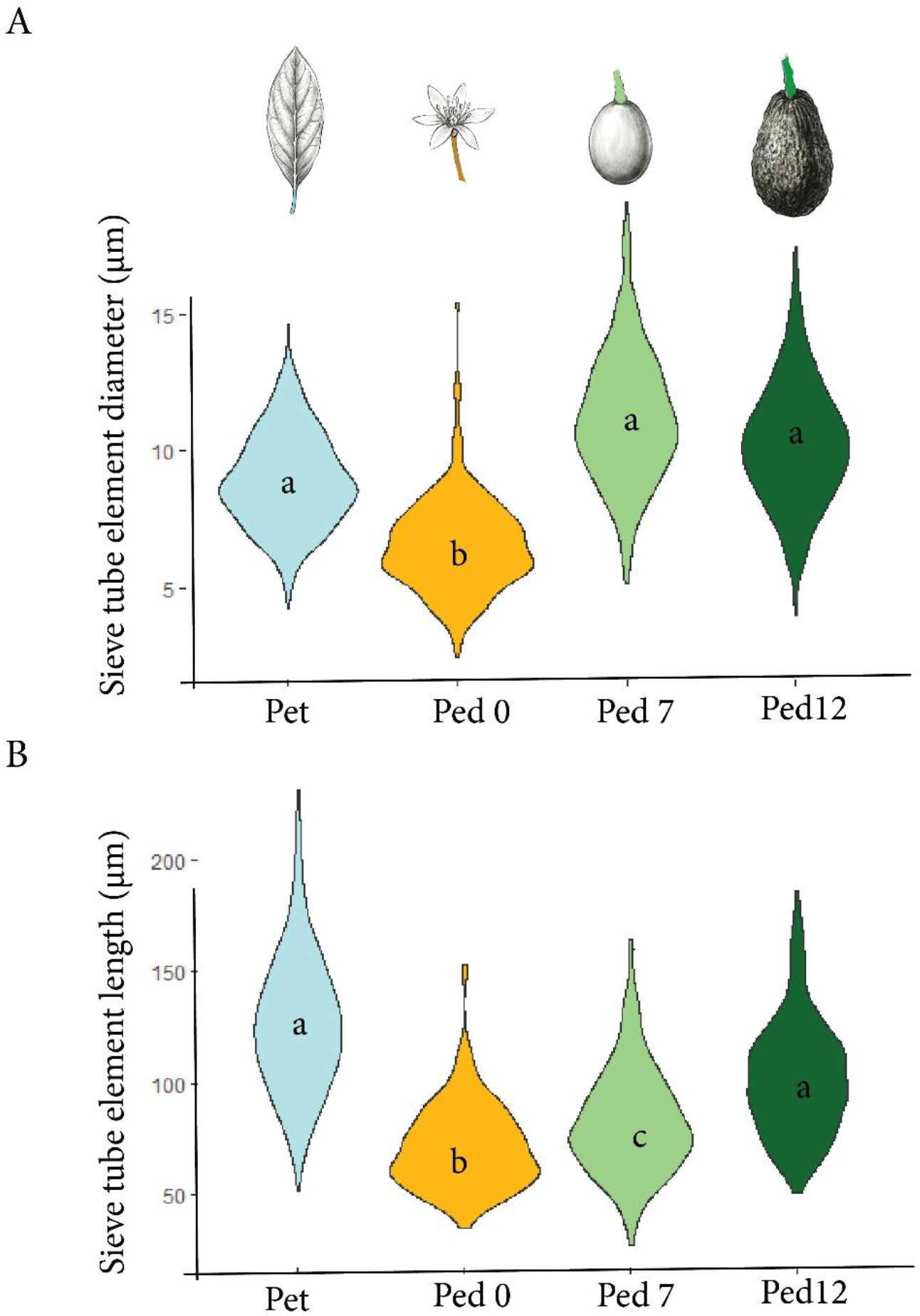
Phloem geometry in the petioles and pedicels of avocado flowers and fruits during development. (A) Sieve tube element diameter. (B) Sieve tube element length. Letters within violin plots indicate significant differences (ANOVA and Tukey’s post hoc test, *P* < 0.05).

#### Sieve plate pore size and number

To understand whether the sieve plate connections followed this pattern (**Figure 6A**), the average radius of the pores connecting the sieve plate was minimum in the flowers (around 0.2µm), doubling this value in developing fruits (0.4µm), but maintaining this size up to the mature fruits (**Figure 6B**). Strikingly, the average number of pores per plate was similar in all the studied developmental stages, without significant differences between flowers, developing and mature fruits, despite the size differences and the prolonged temporal interval of 10 months between both stages (**Figure 6C**). Maintenance of sieve tube element features during fruit development led to quantifying their number.

**Figure 6.**
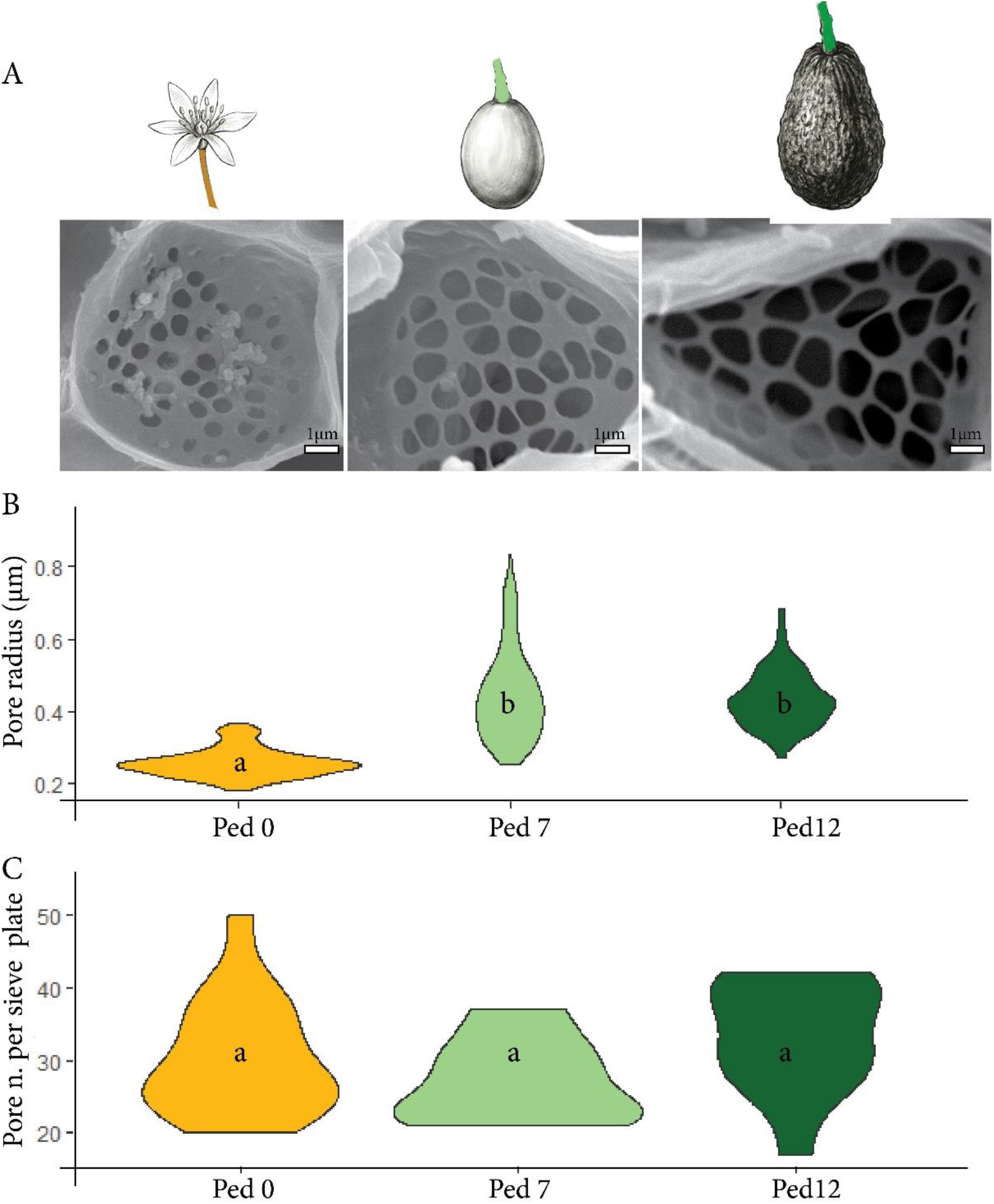
Sieve pore measurements in the pedicels of avocado flowers and fruits during development. (A) SEM images of sieve plates in the pedicels of flowers (Ped 0), developing fruits in July (Ped 7) and mature fruits in December (Ped 12). (B) Average pore radius per plate at each developmental stage. (C) Average number of pores per sieve plate at each developmental stage. Letters within violin plots represent significant differences (ANOVA and Tukeýs post hoc test at *P* <0,05). (A) scale bars: 1µm.

#### Sieve tube element number

Cross sections of the petioles and the pedicels revealed that the size of the axial areas of the phloem correlated directly with the number of sieve tube elements in both peduncles of sources and sinks (including flowers, developing fruits, and mature fruits) (**Figure 7A**). The size of axial areas increased linearly with development, reaching a maximum at the mature stage of the fruits. The number of tubes followed a similar pattern. It changed from ±6,000 tubes in the pedicel of flowers to nearly ±20,000 tubes in mature fruits (**Figure 7B**). Strikingly, the density of sieve tube elements (number per unit area) was maintained constant during all developmental stages, varying between 9,000 and 10,000 per mm^2^.

**Figure 7.**
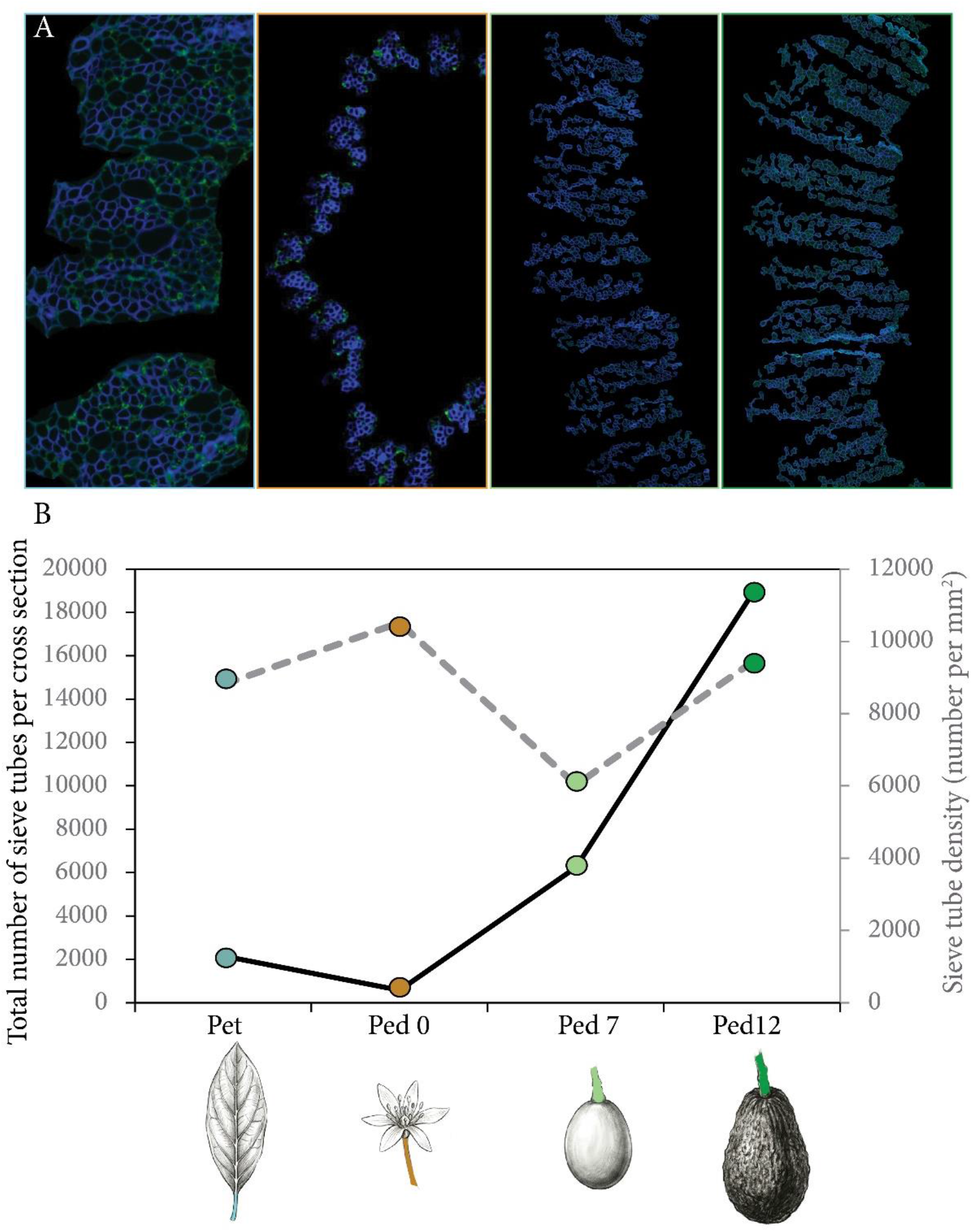
Sieve tube element number per cross-section in petioles and pedicels during development. Tissues analyzed include petioles (Pet) and pedicels of flowers (Ped 0), developing fruits in July (Ped 7), and mature fruits (Ped 12). (A) Processed images stained with calcofluor White for cellulose and pectins (blue color) with isolated phloem at each developmental stage. (B) Total number of tubes per cross-section (left Y-axis) and sieve tube element density (right Y-axis) at each developmental stage.

### Conductivity and conductance in the source and sink organs of avocado

Flow of sugar through the sieve tube elements of the phloem can be described by the pressure-driven flow hypothesis (Horwitz, 1954; Münch 1930; Jensen *et al*., 2013, Knoblauch *et al*., 2016), using as a model the Hagen-Poiseuille equation (Thompson & Holbrook, 2003)

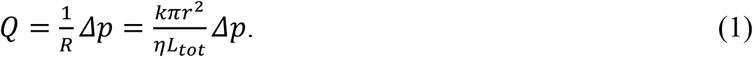

where *Q* is the volumetric flow rate, *Δp* the osmotic pressure difference between source and sink, *R* the hydraulic resistance, *k* the hydraulic conductivity, *η* the viscosity of the fluid, *r* the radius of the total conductive area, and *L*_tot_ the length of the conducting tube. Another measure is the flux, which is invariant of the total cross-sectional area as it is calculated to *q* = *Q*/(*πr*^2^).

Hydraulic conductivity is a widely used measure of fluid flow in the phloem and can be calculated from anatomical data (Thompson & Holbrook 2003b, Mullendore 2010, Jensen 2012b). Following equation (1), the conductivity can be defined as

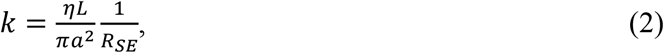

where *R*_SE_ is the hydraulic resistance of one sieve element, *L* the length of the sieve element, and *a* its radius. The hydraulic resistance has been mathematically described before as the sum of the lumen and plate resistance, making use of the Hagen-Poiseuille equation (Jensen 2014).

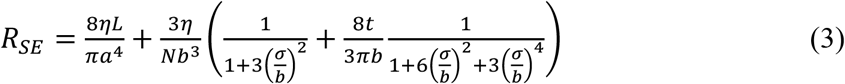

where *t* is the thickness of the sieve plate, and *b* is the average pore radius with the standard deviation *σ*. While the conductivity is specific to one sieve element and can be used to compare different plants, the total conductance *C*, defined as *C* = 1 /*R,* allows us to consider the number of sieve elements per cross-section. Similar to Jensen *et al*. (2012) defining the resistance of one sieve element, we can apply the electrical circuit theory to add multiple sieve elements in one cross-section by adding their resistance in parallel (1/*R* = 1/*R*_1_+1/*R*_2_), and various elements in the longitudinal direction will be added as a serial connection (*R* = *R*_1_ + *R*_2_). Under the assumption that the phloem geometry is the same along the whole translocation length *L*_tot_, the total resistance (*R_3D_*) is

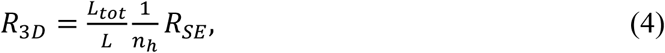

where *n_h_* is the number of sieve elements per cross-section. Similarly, the conductance is

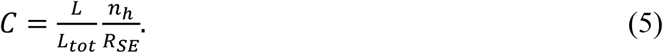

Using eqns. (2) and (3), we calculated the hydraulic conductivity in the avocado leaf as well as in the flower and the fruit (**Figure 8**), where two developmental stages were chosen, month 7 (initial fruit set) and month 12 (mature fruits). Within the veins of an avocado leaf, the conductivity of the sieve elements changes by about 2/3 from 0.3 to 0.45 µm^2^. The conductivity in the flower is much lower than in the leaf, with approximately 0.1 µm^2^ (**Figure 8**). During the development of the fruit, the conductivity increases by 150 % from flower to young fruit at the month 7 of the year. Then it increases further by an additional 60 % to the mature fruit at the month 12, where a hydraulic conductivity of about 0.4 µm^2^ is reached.

**Figure 8.**
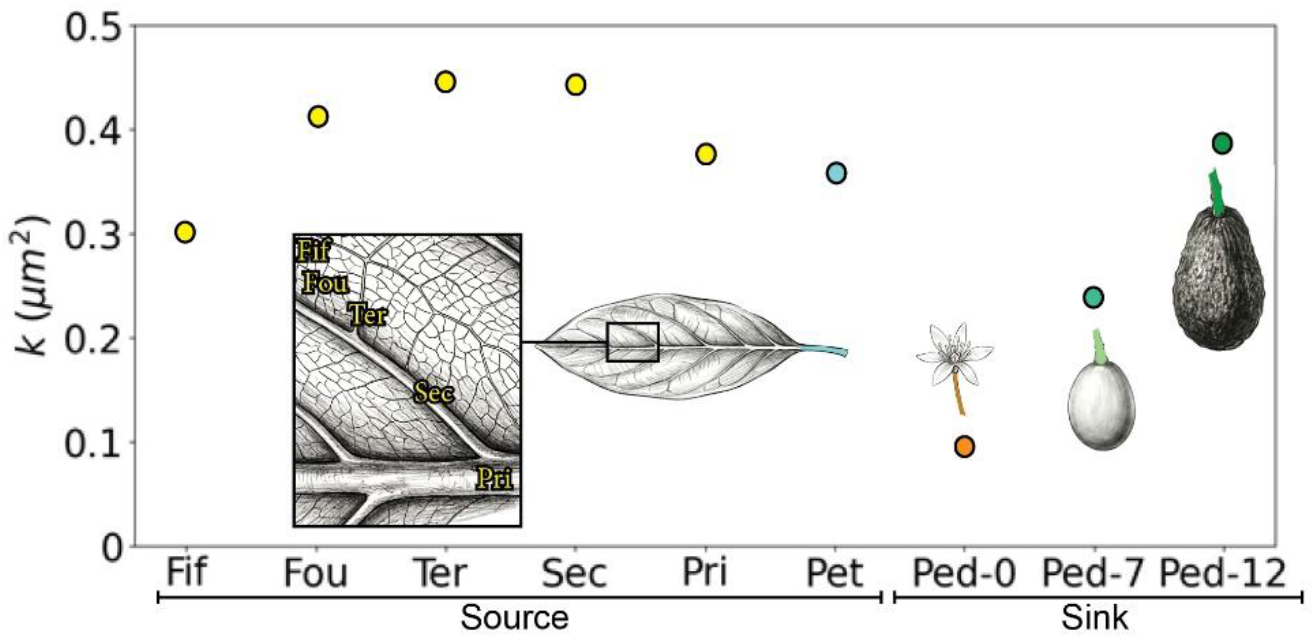
Sieve tube conductivity from source to sink tissues over avocado development. Conductivity was calculated from phloem in source tissues—fifth-order (Fiv), fourth-order (Fou), tertiary (Ter), secondary (Sec), and primary veins (Pri), and petiole (Pet)—to sink tissues, including the pedicels of flowers (Ped 0), developing fruits in July (Ped 7), and mature fruits (Ped 12).

Next to the conductivity, the total phloem conductance (*C_tot_*) was determined at the different developmental stages of the fruit pedicel (**Figure 9A**) using eqn. (5) together with the experimental number of conduits per cross-section (n_h_, see **Figure 7**) and an initial estimation of 1 m for the distance between source and sink (L_tot_). The conductance of the phloem increased from flower to fruit and from developing to mature fruits. This growth is mainly explained by the increase in sieve element number, which scales proportionally with the conductance (eqn. (5)).

**Figure 9.**
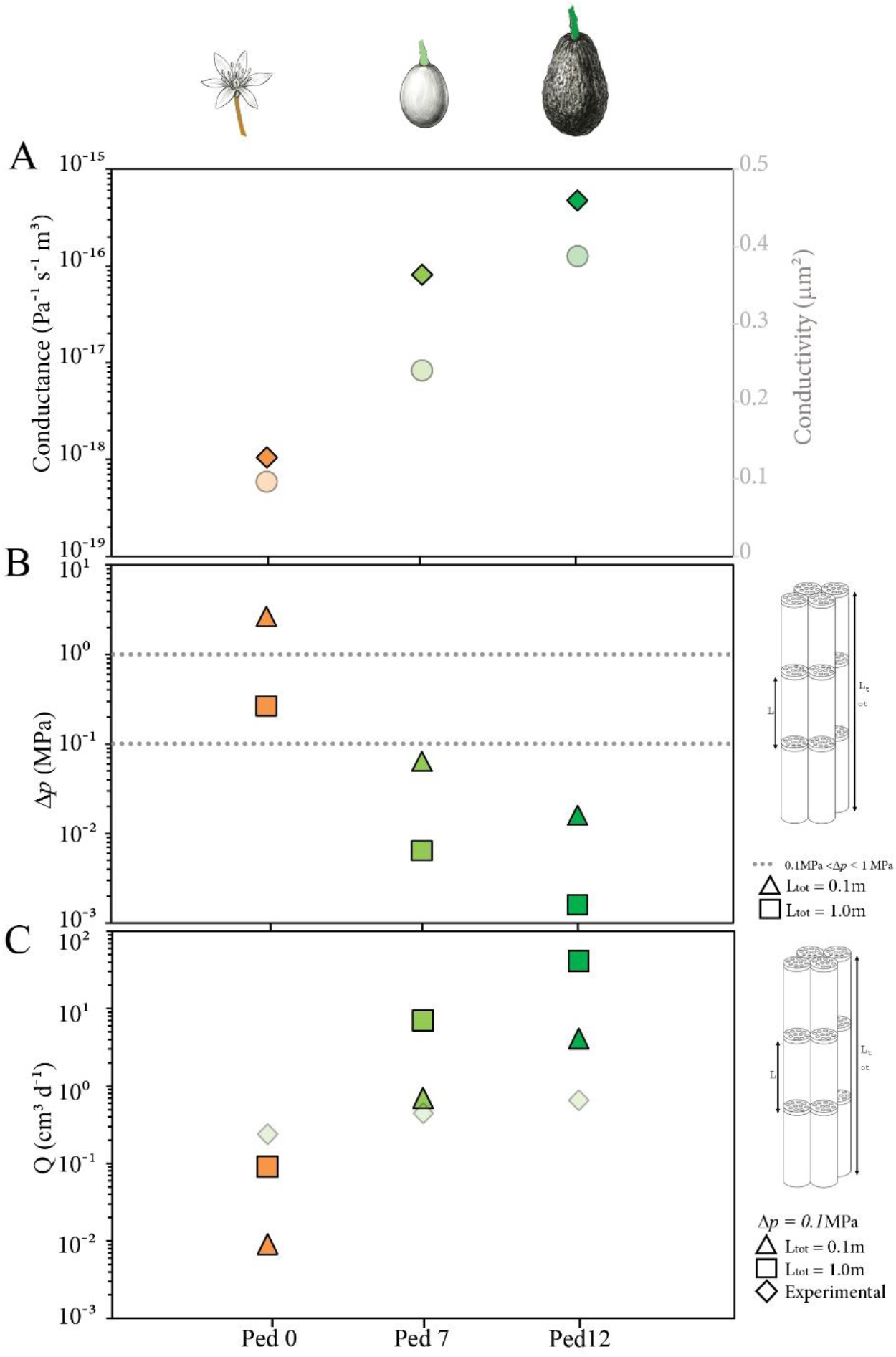
Modeling phloem transport in avocado from source petioles to sink pedicels. Tissues modeled include sink pedicels of flowers (Ped 0. orange), developing fruits (Ped 7, light green), and mature fruits (Ped 12, dark green). (A) Conductance (ribbons) and conductivity (circles) during fruit development show similar trend of increase. (B) Calculated pressure drop of the phloem considering a fixed sap flow (Q) and a predicted phloem operation within the 0.1 m–1.0 m length range (triangles and squares). (C) Calculated flow rate Q considering a fixed pressure drop of 0.1 MPa. within the 0.1 m–1.0 m length range (triangles and squares), compared with experimental values (ribbons).

To verify the calculated values for biological relevance, we further calculated the pressure drop that would result from the conductance (**Figure 9B**). The pressure gradient to drive the viscous flow in a plant is of the order of 0.1-1 MPa (Jensen *et al*., 2013). We therefore expect a pressure gradient within the same range. The relationship between conductance, *C*, and pressure drop, *Δp*, is given by eqn. (1) (keeping in mind that *C* = 1/R).

To estimate the sugar import flow rate *Q,* we rely on measurements of the fruit growth rate *G* (**Supplemental Figure S1**)

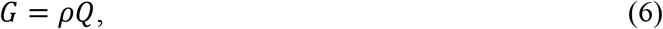

where *ρ* is the density of the phloem sap. Here, we assume a typical sucrose concentration of 18% wt (Jensen *et al*., 2013).

From Eqns. (4) and (5), the pressure gradient can be evaluated as a function of the source-to-sink distance *L_tot_*. . Here, we focus on two examples: *L_tot_* = 0.1m and 1m. The inferred pressure gradients generally fall within or below the typically observed range (0.1-1 MPa, see Jensen *et al*., 2013). While values below 0.1 MPa are not usually observed, they are physically achievable by the plant, unlike pressure gradients above 1 MPa. Variations to the pressure gradients calculated here, could be explained by our general assumption of a uniform system with no change in phloem geometry across the transport area between source and sink. In general, the pressure drop required to achieve the observed growth decreases with increasing fruit maturity, likely due to an increase in the number of sieve tubes per cross-section and, thereby, conductance (**Figure 9B**).

In contrast, we calculated the resulting flow rate at a fixed pressure drop of 0.1 MPa (**Figure 9C**). While the results show a variation between the experimentally estimated flow rates from eqn. (6) to the calculated ones, any differences might be explained by our assumptions of a uniform system; however, both this result and the estimated pressure drop show that a higher flow is theoretically possible.

## Discussion

This empirical study evaluates the role of phloem conduit number from source organs (leaf petioles) to the pedicels of developing fruits over time, addressing a knowledge gap regarding the dynamic nature of phloem architecture in developing sinks. Our findings challenge the idea of continuous scaling of conduit size in fruit pedicels, instead highlighting a bimodal strategy: a rapid increase in individual conduit efficiency during initial fruit set, followed by a sustained rise in total conductive area through phloemogenesis.

### Initial bottleneck during avocado fruit development: adaptation by size

Our work quantified, for the first time in a fruit tree, changes in phloem structure (sieve tube element geometry, sieve plate morphology, and conduit number) in avocado pedicels from flower to mature fruit over time (months). These evaluations revealed a critical initial limitation. In avocado flower stalks, the measured sieve tube elements showed a minimum size (approximately 0.1µm radius) and the smallest pore radii, resulting in a high hydraulic resistance (Jensen *et al*., 2012; Savage *et al*., 2017). This low conductivity coincides with the highest nonstructural carbohydrate concentration (20% of fructose and 35% glucose). Accumulation of sugars within flowers indicates a transient impairment in transport, suggesting hydraulic translocation of photoassimilates through the phloem to the flowers is initially limited (Wen *et al*., 2022; Ren *et al*., 2023). By maximizing solute concentration, the flowers maintain a steep osmotic pressure gradient across the sieve plates, thus driving the viscous flow despite the structural limitation, as predicted by the Münch model (Münch, 1930; Liesche, 2022). Furthermore, a handful of recent studies quantifying phloem transport to floral tissues have reported low conductivity, as in *Paeonia* flowers (Yang *et al*., 2023). The anatomy of the phloem is an essential aspect for understanding both the mechanical properties of the reproductive organs and their greater vulnerability to extremes in the environment compared with leaves (Alves Aun *et al*., 2024). In avocado, a significant carbon budget is implied during the flowering period, given the thousands of flowers per tree in the spring, but most flowers will eventually drop off, given the low fruit set in this species (Alcaraz *et al*., 2021). The high resistance in the floral phloem may serve as a protective mechanism that modulates the initial carbon allocation to reproductive tissues (Turgeon, 2010), leaving a high osmotic pressure within flowers that drives water import during the initial fruit elongation stages.

The 150% increase in sieve tube element conductivity observed from flower to initial fruit set (young fruits at month 7) is a critical developmental adjustment. This is driven by the doubling of the sieve plate pore radius and the tube itself. Given the flow’s proportionality to the fourth power of the radius (Hagen-Poiseuille model), this slight increase represents a massive reduction in resistance, effectively overcoming the initial hydraulic bottleneck (Thompson, 2006). This rapid adjustment establishes an efficient minimum conduit size, allowing the pedicel to meet the rising demand for photosynthates as the fruit begins its rapid growth phase (Falchi *et al*., 2020). Unloading mechanisms in fruits are poorly understood, but previous works in fruit crops such as grapes (Zhang *et al*., 2006) revealed a developmental shift from passive to active phloem loading (Ma *et al*., 2019). While these unloading mechanisms are mainly related to physical gates of phloem cells in fruits (i.e. plasmodesmata or active transporters such as sucrose transporters), the role of phloem structure and its consequent hydraulics has been mostly overlooked. Our results point to a developmental hydraulic shift from a highly resistant phloem during flowering to a less resistant one during the initial stages of fruit development, depicted by an increase in the geometrical parameters of sieve tubes in the pedicel of fruits that achieved a minimum size.

### Sustained growth of avocados during fruit elongation: adaptation by number

Following the fruit set, there was a maintained growth of fruits until month 10, but the pedicel showed a constant diameter. Dry matter accumulation over time, coincident with the continuous drop of carbohydrates in the developing fruits, demonstrates that carbohydrate transport is consistent with an enhanced phloem conductivity (Thompson, 2006). This is in line with previous evaluations of starch accumulation in the stems of avocado, which showed a similar trend during development (Whiley *et al*., 1996; Liu *et al*., 1999). While water transported through the xylem is required for cell elongation during fruit growth (Marcelis and Heuvelink 2007; Zamski 2017), and the import/export flow rates of water have been previously measured in avocado (Kaneko *et al*., 2024), the structural capacity of the phloem to support the required carbohydrate transport rates has been traditionally ignored.

Our findings show that after the initial geometric change (month 7), the length, width, and sieve plate characteristics remain statistically constant until month 12. In contrast, total phloem conductance (*C*) increases monotonically throughout fruit development, closely mirroring the continuous increase in the number of sieve tubes (from 6000 to 20000 per fruit pedicel). The maintenance of a constant sieve tube density (approximately 8000 to 10000 conduits per mm2) strongly suggests the plant’s strategy is to increase the total cross-sectional area of the phloem through cambial activity in the pedicel, a process termed phloemogenesis. Previous reports with giant pumpkins strongly suggested that fruit size was strongly related with the capacity of phloem transport (Savage *et al*., 2015), and the phloem variations with seasons has been mainly studied in temperate species, given the shifts that accompany the formation of new leaves and fruits during the spring (Ray and Savage, 2021). The seasonal effect has been less explored in evergreen species such as avocado, but their prolonged fructification periods allow to understand the role of biosynthesis of sieve tube elements over time as critical for a proper maturation and, therefore, an efficient crop harvest.

This is a strategy of scaling-by-number rather than scaling-by-size. Interestingly, this approach is different from source organs (leaves), where the STE size scales with vein order (Carvalho *et al*., 2018; Losada & Holbrook, 2019; Losada *et al*., 2022). This fundamental difference in vascular developmental regulation ensures that total conductive capacity keeps pace with the fruit’s mass accumulation rate, preventing the phloem from becoming a limiting factor. The continuous structural differentiation reflects a dedicated resource investment to maintain the transport capacity required by a high demand sink (Stanfield *et al*., 2024). The scaling-by-number strategy aligns with our hydraulic models suggesting that increasing conduit number is highly effective for boosting long distance transport (Liu *et al*., 2024). The linear rate of fruit growth thus determines the capacity of phloem transport, suggesting a necessary synchrony driven by the sink. The continuous structural differentiation implies that sink tissues require a degree of plasticity (Jensen, 2018), in contrast to the more static vascular tissues of mature source organs.

### Conclusions and future directions

Our data demonstrates that the hydraulic transport capacity of the phloem in avocado pedicels adapts to increasing demands through a staged mechanism: initial size optimization, followed by a sustained scaling-by-number (phloemogenesis). This finding provides essential anatomical data necessary for refining whole plant carbon allocation models (Sevanto, 2018).

The question remains on the internal and external regulation of this phloemogenesis, and whether this strategy could be generalized to other fruits with prolonged development. Future research should focus on the regulatory signals that activate this sustained phloemogenesis in the pedicel, investigating the role of hormonal gradients (e.g. auxin and gibberellin, which are known to influence cambial activity, and turgor pressure feedback from the fruit, which has been implicated in sinks like japanese plum (Grappadeli *et al*., 2019). Further comparative studies are needed to determine if this bimodal scaling strategy is universal among economically important, highly demanding fleshy fruits, necessary to understand developmental mechanisms governing reproductive development.

## Methods

### Plant material, size, and relative water content of leaves, flowers, and fruits

A subset of 41 trees (3-4m tall) belonging to an orchard of avocados cv’Has’, >20 years old, planted at 8 x 8m, irrigated on demand, that belongs to the germplasm collection of the Institute for Mediterranean and Subtropical Horticulture ‘La Mayora’ – CSIC -UMA (4°2ʹ24.32ʹʹN, 36°45ʹ32.75ʹʹE), which is located in the subtropical microenvironment of Málaga province in southern Spain.

Three developmental stages of reproductive organs were selected to evaluate: flowers, fruits after the first fruit drop -July, month 7-, fruits at the maximum size -October, month 10-, and mature fruits for harvest -December, month 12-. The diameter of the pedicel sustaining developing fruits was measured with a caliper, along with the fruit’s maximum length and diameter. Data on the diameter of the flower pedicels and carpel size from flowers were obtained from a previous work carried out in the same orchard (Alcaraz *et al*., 2010).

To calculate the relative water content (RWC), we collected three leaves per tree from a total of 10 trees (n=30), 15 belonging to previous year leaves (December), and 15 belonging to newly expanded leaves (may), 11 groups of 10 flowers each (n=110 flowers), 11 fruits from month 7, and 11 fruits from month 12. They were all weighed fresh (Fw), except for the small flowers, which were weighed in groups of 10 on a four-decimal scale. All samples were dried in the oven at 60°C for at least 48 h and weighed thereafter to get the dry weight (Dw). Relative water content per sample was calculated following the protocol of Poorter *et al*. (2009). In parallel, we measured dry matter content using the non-invasive near-infrared (NIR) method (F-751, Felix Instruments) monthly from the first fruit drop (month 7) to maturity (month 12). Note that the small size of the flowers makes it impossible to measure their dry matter using this method (**Figure**S1).

### Non-structural carbohydrate evaluation

Fully developed leaves from the previous year (i.e. when the fruit of the current year are mature or month 12) and fully expanded leaves from the current (i.e. at initial fruit set or month 7), were cut from 15 adult trees (3 leaves per tree, n=45) and used along with a subset of 10 flowers per each tree (n=150), two fruits per tree at the initial fruit set (n=30), and two fruits per tree at the mature stage (n=30). Immediately after collection, the material was incubated in an oven at 100°C for 1 hour to deactivate enzymes, and then the samples were dehydrated in the same oven at 70°C for 3 days. Then, samples were homogenized to a fine powder using a ball mill (TissueLyser, by Qiagen Inc., Mississauga, ON, Canada).

For the quantification of non-structural carbohydrates (NSCs), a modified protocol of Landhäusser *et al* (2008) was followed. First, 30 mg of powder from each sample was weighed, and NSCs were extracted using 80% ethanol. The enzymatic assay and absorbance readings were performed in 96-well polystyrene plates (Thermo Scientific™ Nunc MicroWell 96-Well Microplates) using a microplate reader (BioTek Instruments, PowerWave XS2, USA). Glucose was directly quantified by an enzymatic assay in a 96-well plate mediated by hexokinase-glucose 6-phosphate dehydrogenase (Sigma G3293) and linked to the production of NADH (absorbance 340 nm). The NADH produced by hexokinase-glucose 6-phosphate dehydrogenase, after 1 hour and 40 minutes of incubation, is directly proportional to the glucose present in the sample. The remaining NSCs were converted by the enzymatic method into glucose, starting by starch digestion using α-amylase from *Bacillus licheniformis* (Sigma A4551) and amyloglucosidase from *Aspergillus niger* (Sigma Cat. ROAMYGLL); the aqueous phase with the hydrolysed glucose was loaded into the 96-well plate along with hexokinase-glucose 6-phosphate dehydrogenase, incubated for 1 hour 40 minutes, and finally, absorbance was measured at 340 nm. Sucrose was first digested in a 96-well plate with an invertase enzyme from baker’s yeast (Sigma I9274) to obtain glucose and fructose. After digestion, each sample was loaded into a new 96-well plate containing hexokinase-glucose-6-phosphate dehydrogenase, incubated, and measured as described above. Then, fructose quantification was performed in a 96-well plate using a combination of phosphoglucose isomerase (Sigma P538) and hexokinase-glucose-6-phosphate dehydrogenase (absorbance at 340 nm). All calculations were performed on a dry-weight basis.

### Fluorescence microscopy for phloem evaluations

To evaluate the size of sieve tube elements, longitudinal sections of vascular elements (leaf veins, petioles, and flower and fruit pedicels) were fresh cut using a micro-scalpel in 1xPBS with Leica S6D Stereo Zoom Microscope, resulting in approximately 1mm sections, which were stained onto a microscope slide with 0.1% aniline blue in PO_4_K_3_ (Currier and Strugger, 1956) to stain callose and 0.1% calcofluor white in 10mM KCl (Hughes and McCully, 1975) to label cellulose. We measured a minimum of 500 tubes per leaf vein, except for minor veins, which counted 266 tubes. The samples were extracted from 10 leaves belonging to different trees, 10 flower pedicels (301 tubes), 10 fruit pedicels from month 7 (306 tubes), and 10 fruit pedicels from month 12 (400 tubes).

This procedure was also applied to transverse sections of the pedicel belonging to 10 mature fruits, which were used to count the number of vascular elements per axial area of the phloem. The number of axial regions was further counted and measured individually using image analysis. Calcofluor white identified the sieve tube elements of the phloem in sections of avocado, and to confirm that counts were indeed tubes, we immunolocalized the walls of sieve tube elements with the LM26 monoclonal antibody (Torode *et al*., 2018). To this end, pedicels and petioles were frozen in liquid nitrogen, then fixed in super chilled 100% ethanol to maintain their integrity, then included in Technovit 8100 resin for several days or weeks, and blocks were made under anoxic conditions at 4°C. Sections of 4µm thickness were obtained with Leica EM UC7 ultramicrotome (Leica Microsystems). Immunolocalization was performed as described before (Torode *et al*., 2018). While the total number of tubes was counted for most developmental stages, the large size of the fruits implied that, after counting the number of tubes in a subset of axial areas, we estimated the total number based on the average number of tubes per area multiplied by the average number of the total regions counted.

The slides stained and immunolocalized were observed under Leica DM LB2 epifluorescence microscope using a 405-nm filter (aniline blue and calcofluor) and a 488nm filter for FITC belonging to immunolocalization. The microscope is equipped with a camera and LAS V4.5 software.

### Scanning electron microscopy for sieve plate quantifications

Small sections of petioles and pedicels from flowers and fruits at months 7 and 12 were cut, immediately submerged in liquid nitrogen to prevent callose accumulation in the sieve pores, and transferred to super-chilled 100% ethanol. The sections were incubated first at −80°C for 24h, and then gradually moved to −20°C for 24h, 4ªC for 24h, and room temperature. 1-2mm sectioned samples were cut on ethanol using a double-edged razor blade. These sections were washed three times in distilled water to remove ethanol. A 0.1% proteinase K solution dissolved in 50mM Tris-HCl buffer, 1.5mM Ca^2+^ acetate, and 8% Triton X-100, pH 8 (Mullendore *et al*., 2010) in a water bath at 60°C for at least 14 days, replacing the mixture once a week. After 2 weeks, the cytoplasmic content was digested, sections were incubated in ethanol to inactivate the enzyme, and then rinsed three times in distilled water. These sections were incubated at 60°C for 24h in a 0.1% α-amylase solution, which removes the starch accumulated on the sieve plates. We rinsed the sections in ethanol, then 3 times in water, and finally removed all the water. Lyophilization was done with a freeze-drier (CoolSafe 4-15L Freeze Dryers, LaboGene, Allerod, Denmark) for 24h. Samples were mounted on SEM studs, covered with gold-palladium using a sputter coater (QUORUM Q 150R ES), and finally observed using scanning electron microscopy (JEOL JSM-840). We measured at least 140 pores from each pedicel of the fruits harvested in months 7 and 12, as well as from flowers.

### Image and data analysis

All microscopic images were processed with IMAGEJ (National Institutes of Health, https://imagej.nih.gov/ij/). Specifically, we used it to measure the length and diameter of sieve tube elements, count sieve plates, and measure the area of each sieve plate/pores. Data analysis was performed using R (version 4.2.1). All data followed a normal distribution, as determined by a Shapiro test. ANOVA of 1 and 2 factors and Tukey’s test at a *p*<0.05 were used to test anatomical variation within pedicels, petioles, fruit size, and sieve pores.

