## Supplementary figure 1 for "Phloem conductance from leaf to fruits in avocado during development"

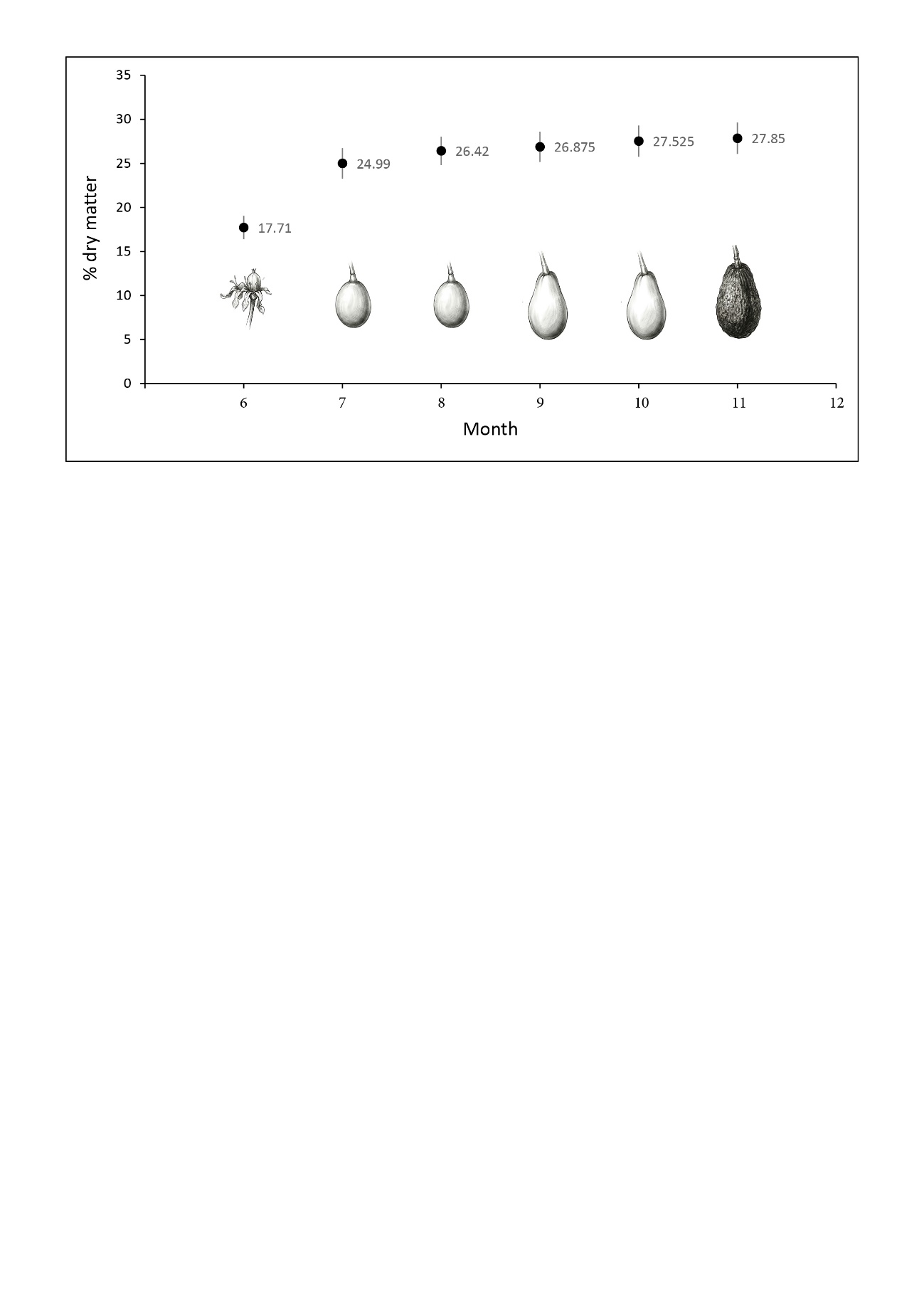
**SUPPLEMENTARY FIGURES**

**Supplementary Figure 1. Dry matter content during avocado fruit development.** Y axis represents the average % of dry matter evaluated with the near infra red methodology (NIR) method in the field. X axis represents the month of the year (6, 7: July; 8: August; 9: September; 10: October; 11: November; 12: December). ­­­­­

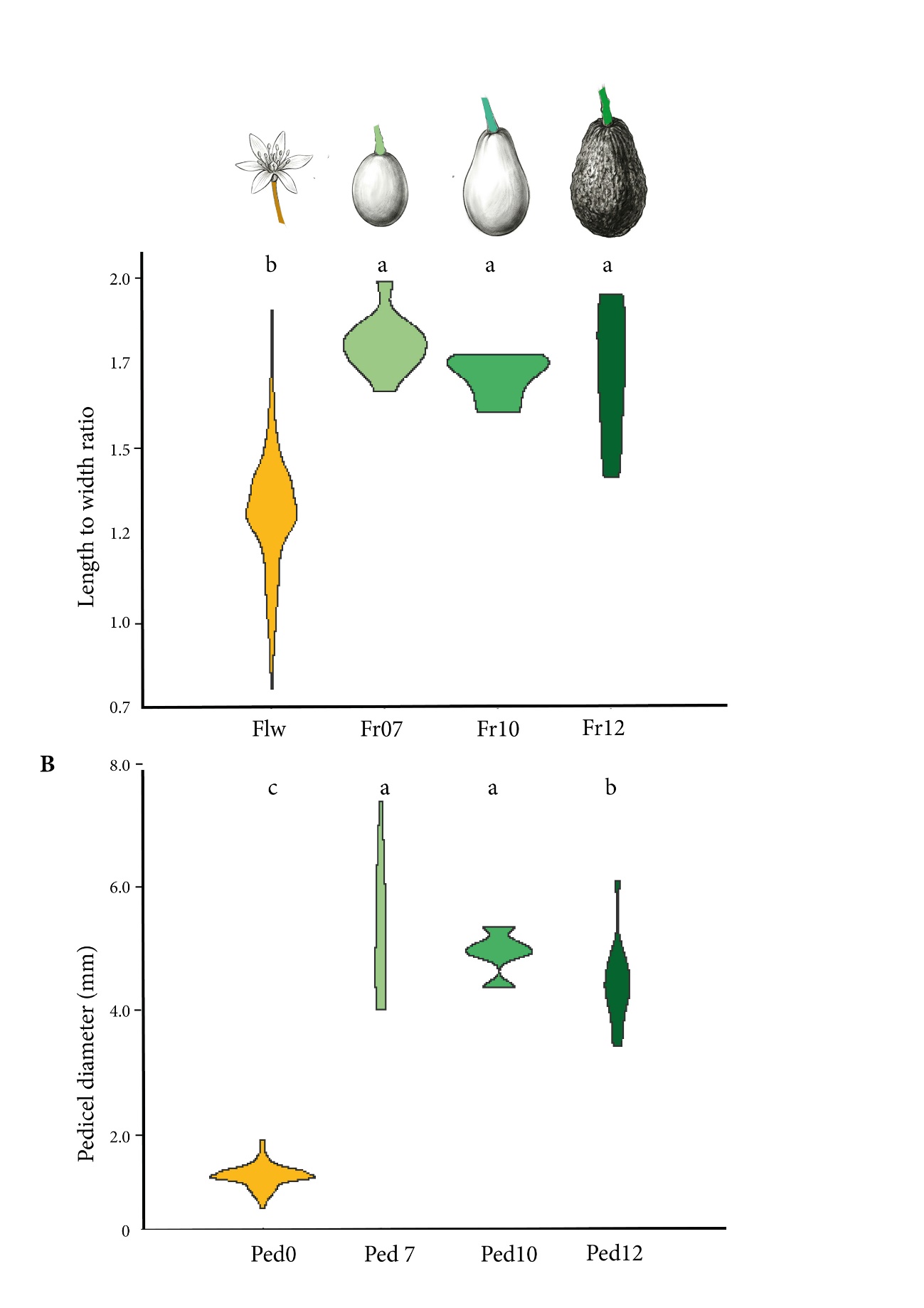

**Supplementary Figure 2. Sink tissue ratio and pedicel diameter during avocado development.** (A) Length-to-width ratio of the flower ovary (Flw) and developing fruits in July (Fr07), October (Fr10), and December (Fr12). (B) Pedicel diameter at the corresponding developmental stages (Ped 0, Ped 7, Ped 10, and Ped 12, respectively). Letters over plots indicate significant differences (ANOVA and Tukey's post hoc test, *P* < 0.05).
